# Sparse Autoencoders Reveal Structural and Family-level Features in BiRNA-BERT

**DOI:** 10.64898/2026.08.11.744228

**Authors:** Mohammad Sadat Hossain, Md. Roqunuzzaman Sojib, Md Toki Tahmid, M Saifur Rahman

## Abstract

**Motivation:** RNA language models learn representations that support structure and function prediction, but which biological concepts their hidden states encode remains unclear. Sparse autoencoders (SAEs) decompose hidden states into interpretable features, yet have not been applied to RNA language models, where byte-pair tokenization breaks the one-token-one-nucleotide correspondence that nucleotide-level attribution assumes.

**Results:** We present SPIRAL, a layer-wise SAE analysis of BiRNA-BERT. Independent SAEs at layers 0, 5, and 11 expand each 768-dimensional hidden state into 6,144 features while preserving model behaviour (explained variance above 0.99997; masked-language-model sequence recovery near 99.7%). Tokenizer-aware offset propagation aligns features to nucleotides: at layer 5, 44.3% of tested features are significantly associated with bpRNA secondary-structure classes (mean enrichment 1.61 *×*), and all 1,237 eligible features with RNAcentral RNA types. Sparse profiles raise *k*-nearest-neighbour balanced accuracy from 0.328 to 0.359 over dense embeddings at layer 5.

**Availability and Implementation:** Source code is available at https://github.com/SadatHossain01/SPIRAL; the code, evaluation data, and trained SAE checkpoints are archived at https://doi.org/10.5281/zenodo.21891845.

**Supplementary information:** Supplementary data are presented alongside the manuscript.

## 1. Introduction

RNA molecules perform a diverse set of cellular roles, including translation, post-transcriptional regulation, and catalysis. These functions are determined not only by the primary sequence but also by secondary structure and family-specific sequence motifs. The rapid growth of RNA sequencing data has led to the development of several RNA language models that learn rich contextual representations directly from nucleotide sequences. Large-scale RNA language models such as RiNALMo (Penić et al., 2025), RNA-FM (Chen et al., 2022) and BiRNA-BERT (Tahmid et al., 2025) now achieve strong benchmark performance on various diverse downstream tasks.

Despite achieving state-of-the-art performance on a range of downstream tasks, RNA language models remain black boxes in terms of understanding what these models actually learn. Decoding hidden representations into biologically relevant features is crucial for assessing model reliability in out-of-distribution settings, interpreting unexpected predictions, and potentially steering model outputs. The fundamental obstacle is the superposition hypothesis (Elhage et al., 2022). A neural network with a fixed-dimensional hidden state must simultaneously represent many more concepts than it has dimensions, so individual hidden-state coordinates inevitably mix multiple unrelated factors. Thus, identifying a feature representing a stem nucleotide or specific RNA family sequence requires methods that can disentangle dense hidden states into more interpretable components.

Sparse autoencoders (SAEs) offer a principled approach to this disentanglement for mechanistic interpretability (Huben et al., 2024; Bricken et al., 2023). An SAE maps each hidden state into a much larger overcomplete dictionary while enforcing an *L*_1_ sparsity penalty that encourages only a small subset of features to be active for any given token. If the reconstruction of the original hidden state remains faithful, each active feature can serve as more interpretable units of analysis than individual hidden-state dimensions, which are often polysemantic. Mechanistic interpretability research in large language models has demonstrated that sparse features trained this way can correspond to coherent semantic properties and support causal interventions (Bricken et al., 2023; Templeton et al., 2024). This makes SAEs a promising framework for asking whether biological concepts encoded by RNA language models can be isolated as individual sparse features.

Recent work has begun applying sparse autoencoders to biological sequence models. SAE-RNA (Kim and Nam, 2025) showed that sparse features from RINALMO correspond to biologically meaningful concepts, including secondary-structure motifs and RNA family-specific patterns. In parallel, sparse autoencoder studies of protein language models have demonstrated mechanistic feature interventions at the amino-acid level (Simon and Zou, 2025; Garcia and ansuini, 2025). However, several questions remain open for RNA models. RNA language models that employ adaptive BPE tokenisation introduce an additional alignment challenge: a single token may span multiple nucleotides. Consequently, nucleotide-level annotations cannot be directly compared with token-level sparse activations and require projection through the tokenizer’s token-to-sequence alignment. While token-to-input alignment is well understood in NLP, it has received comparatively little attention in RNA interpretability, where nucleotide-resolution analyses are often required. Existing RNA SAE analysis has primarily analyzed a single layer output, without exploring how such features change across the transformer depth.

We present SPIRAL (**Sp**arse autoencoders for **I**nterpretable **R**NA **A**na**l**ysis), a layer-wise SAE analysis of BiRNA-BERT, an RNA language model designed with an adaptive byte-pair tokenization scheme for representing local as well as long-range structures of RNA. In BPE tokenization, a single model token may span multiple nucleotides, whereas many biological annotations, including secondary-structure labels, are defined at nucleotide resolution. SPIRAL addresses this challenge by combining faithful SAE reconstruction, tokenizer-aware nucleotide alignment, and biological validation across both structure-level and family-level tasks. Our main contributions are as follows.

1. **Trained SAE dictionaries at three representative layers**. We train SAEs at layers 0, 5 and 11 of BiRNA-BERT (6,144 features, 8*×* expansion) and observe different outcomes across layers, motivating layer-specific tuning. We evaluate each SAE not only by hidden-state reconstruction, but also by preservation of masked-language-model outputs and empirical sparsity.
2. **Nucleotide-resolution structure alignment via BPE offset propagation**. We introduce BPE offset propagation to map token-level sparse activations to nucleotide-level bpRNA structure labels. This reveals layer-dependent structure selectivity, with middle layers containing enriched structural contexts.
3. **RNA-type alignment and representation utility**. We evaluate sparse profiles on a 16-type RNAcentral holdout and show that the layer-5 SAE improves *k*NN RNA-type classification.

## 2. Methods

### 2.1. Model, Data, and Sparse Autoencoders

SPIRAL studies a frozen BiRNA-BERT encoder loaded from the public weights^1^ (Tahmid et al., 2025). BiRNA-BERT is a 12-layer transformer with 768-dimensional hidden states and adaptive BPE tokenisation. Because one BPE token may cover multiple nucleotides, all nucleotide-level analyses use the tokeniser offset map rather than assuming a one-token-to-one-nucleotide alignment. We extract hidden states from layers 0, 5, and 11, representing the input-proximal, middle, and final encoder layers.

The sparse autoencoders are the only models trained in SPIRAL. For each analysed layer, we train an independent SAE on non-special-token activations from the first 999,744 RNAcentral sequences^2^ (The RNAcentral Consortium, 2019). Hidden states are standardised per layer using a fixed activation mean and standard deviation estimated from the first 100,000 training sequences.

For evaluation, the nucleotide-level secondary-structure analysis uses 25,000 bpRNA-90 sequences^3^ (Danaee et al., 2018) with nucleotide-level structure annotations. RNA-type analysis uses a separate RNAcentral holdout of 25,000 sequences spanning 16 RNA types. Because RNAcentral is also the SAE training source, we compared post-training-window candidates against the one-million-sequence training reference using cd-hit-est-2d at 80% global sequence identity (word size 5), as found in literature (Tong and Liu, 2019; Singh et al., 2019; Qiu, 2023). Sequences were truncated to at most 510 nucleotides, and required to contain at least 10 nucleotides. Further information about the construction process is provided in Supplementary Table S1.

For each layer, the SAE maps a standardised BiRNA-BERT hidden state **x** ∈ ℝ^768^ to an overcomplete 6,144-dimensional sparse feature vector and reconstructs it in the same standardised space:

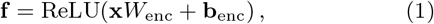

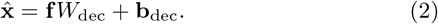

Here **f** is the learned sparse representation. The training objective balances reconstruction fidelity with sparsity:

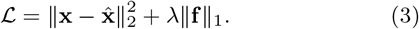

The reconstruction term forces the SAE to preserve information present in BiRNA-BERT, while the *L*_1_ term discourages using many dictionary features at once. This trade-off is the reason SAEs are useful here: if reconstruction is accurate but only a small subset of features is active for a token, individual features become easier to inspect biologically than dense hidden-state coordinates. A feature is treated as active for a token when *f*_*i*_ *>* 0. All SAEs use an 8 *×* expansion factor, *L*_1_ coefficient *λ* = 10^−3^, ReLU encoder activations, untied encoder and decoder weights, AdamW with cosine learning-rate decay, five training epochs, a batch size of 1,024 token activations, learning rate 10^−3^, weight decay 10^−4^, and gradient-norm clipping at 1.0. Encoder weights used Kaiming initialization (He et al., 2015) and decoder weights used Xavier initialization (Glorot and Bengio, 2010); no decoder unit-norm constraint was applied. The training configuration checked for dead features every 50,000 optimization steps using a 10^−5^ activation frequency threshold over 50 sampled batches and resampled features below that threshold.

### 2.2. Reconstruction and Sparsity Evaluation

We evaluate the sparse autoencoder representation along three major axes: hidden-state reconstruction, preservation of masked-language-model outputs, and sparsity of the learned feature code. A single metric is not enough: a reconstruction can be geometrically close to the original representation while changing the language-model output, or it can reconstruct well only by activating so many features that the representation is no longer interpretable. We therefore evaluate complementary properties.

Explained variance quantifies how much hidden-state variation survives the SAE bottleneck:

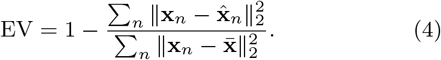

Values near 1 indicate that very little variance is lost. Mean cosine similarity checks whether original and reconstructed hidden states point in the same direction,

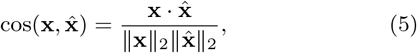

which matters because downstream neural computations often depend on direction as well as magnitude. To verify that reconstruction fidelity also holds at the model-output level, we pass reconstructed hidden states through the masked-language-model head. Sequence recovery records the fraction of positions for which the top predicted token remains unchanged, KL divergence compares the full token-probability distributions between the original output distribution (P) and the SAE-reconstructed output distribution (Q),

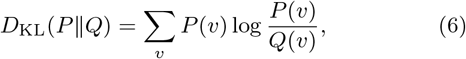

and cross-entropy change ΔCE = CE_SAE_ − CE_base_ shows whether reconstruction makes the original masked-token task easier or harder. Finally, empirical sparsity is measured by mean *L*_0_,

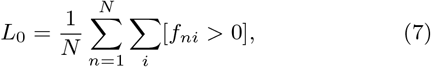

the average number of active SAE features per token. Here, a feature is counted as active when its activation is greater than zero. This is the interpretability guardrail: high fidelity with low-to-moderate *L*_0_ suggests a compact decomposition, whereas high fidelity with very dense activations would be closer to copying the original representation.

### 2.3. Secondary-Structure Alignment

Secondary structure labels were obtained from the bpRNA-90 dataset (Danaee et al., 2018). The Pseudoknot (K) class was excluded because it is absent from the evaluated subset. Consequently, our analysis considered the remaining seven secondary structure classes: External loop (E), Stem (S), Hairpin (H), Internal loop (I), Multi-loop (M), Ambiguous (X), and Bulge (B).

Since a BPE token can span multiple nucleotides, activations are aligned using offset mappings and replicated across all covered positions. For a BPE token *t* with nucleotide span [*s*_*t*_, *e*_*t*_), an active SAE feature is credited to every nucleotide covered by that token. For the count for feature *i* and structure label *ℓ* pair (*i, ℓ*), we sum the number of nucleotides labeled *ℓ* within every token whose activation for feature *i* is greater than zero.

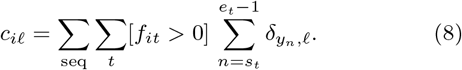

Here, 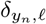 is the Kronecker delta, equal to 1 when nucleotide *n* has structure label *ℓ* and 0 otherwise. This offset propagation preserves the biological span of multi-nucleotide BPE tokens instead of forcing each token onto a single arbitrary nucleotide. The structure analysis then asks three linked questions.

First, does the feature exhibit an activation pattern across labels that is unlikely to arise under the bpRNA background distribution? We use Pearson’s *χ*^2^ test to quantify how significantly the observed activations diverge from an expected random background distribution,

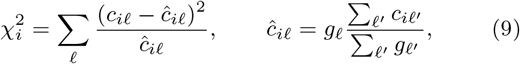

where *g*_*ℓ*_ is the global nucleotide-level background count for label *ℓ*, and the expected count *ĉ*_*iℓ*_ assumes the feature activates proportionally to the dataset’s base rates. Bonferroni correction is applied across tested features. We use this test to define the significant-feature counts reported in Table 2; effect magnitude is summarized separately by selectivity and enrichment.

**Table 1.** Reconstruction quality of sparse autoencoders (SAEs) trained at BiRNA-BERT layers 0, 5, and 11. Explained variance and cosine similarity measure hidden-state fidelity; sequence recovery, KL divergence, and CE Δ measure preservation of masked-language-model output; mean *L*_0_ is the average number of active features per token, with the percentage of active dictionary features shown in parentheses. Lower KL and mean *L*_0_ are better, and CE Δ values closer to zero indicate better output preservation.

| Layer | Explained variance | Cosine similarity | Sequence recovery | KL divergence | Mean $L_0$ (%) | Cross Entropy $\Delta$ |
| --- | --- | --- | --- | --- | --- | --- |
| 0 | **0.999989** | **0.999996** | **0.9977** | $1.01 \times 10^{-5}$ | **975 (15.9)** | **+0.000318** |
| 5 | 0.999980 | 0.999992 | 0.9972 | $1.04 \times 10^{-5}$ | **775 (12.6)** | +0.001414 |
| 11 | 0.999976 | 0.999991 | 0.9970 | $2.14 \times 10^{-5}$ | 791 (12.9) | +0.000961 |

**Table 2.** Secondary-structure alignment summary across BiRNA-BERT layers. “Tested” counts features active in at least five sequences; “Significant” counts features passing a Bonferroni-corrected Pearson *χ*^2^ test against bpRNA-90 nucleotide secondary-structure labels. Median selectivity is the median fraction of activations on the preferred label, and mean enrichment is preference relative to background frequency, averaged over all tested features.

| Layer | Tested | Significant | Median sel. | Mean enrich. |
| --- | --- | --- | --- | --- |
| 0 | **6,139** | **5,988(97.5%)** | 0.453 | 1.04× |
| 5 | 4,053 | 1,794(44.3%) | **0.464** | **1.61×** |
| 11 | 5,483 | 3,980(72.6%) | 0.463 | 1.21× |

Second, we ask how pure the feature’s preference is. We quantify structural preference with a selectivity score defined as the fraction of activations assigned to the feature’s most frequent structure:

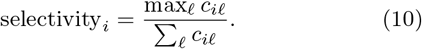

A value near 1 means that nearly all activations fall on one structure class, the uniform-class reference is 1*/*7.

Third, is the preferred structure more common for this feature than expected from the background? Metrics like raw selectivity fail to account for baseline structure frequencies, common structures can appear artificially favored even when activations are random. For the preferred label *ℓ*^∗^ = arg max_*ℓ*_ *c*_*iℓ*_, we compute

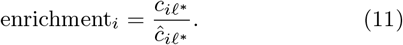

Selectivity and enrichment are interpreted together. Selectivity reflects the categorical purity of a feature, while enrichment serves as protection from the majority class bias, making sure that the feature is truly specialized and not reflective of the underlying background distribution.

### 2.4. RNA-Type Alignment and Representation Utility

RNA-type analyses use the similarity-filtered RNAcentral holdout described above. This holdout contains 25,000 sequences spanning 16 RNA types and is used only after SAE training is complete.

For each sequence *q*, token-level SAE activations are mean-pooled into a sequence profile,

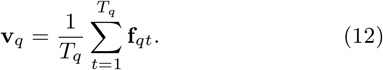

where *Tq* denotes the number of tokens in sequence *q* and **f**_*qt*_ is the SAE activation vector at token *t*. The resulting profile **v**_*q*_ represents the average activation strength of each SAE feature across the entire sequence.

Only features whose sequence-mean activation exceeds 0.05 in at least 10 sequences are tested. For each eligible feature dimension, we compare activation distributions across the 16 RNA types with the Kruskal-Wallis H-test. We use Kruskal-Wallis rather than ANOVA because SAE activations are sparse, non-negative, and heavy-tailed. Its test statistic is

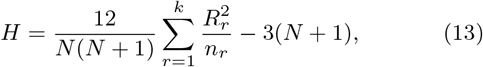

where *R*_*r*_ is the sum of ranks in RNA type *r, n*_*r*_ is that type’s sample size, *k* = 16, and *N* = 25,000. A significant *p*-value says that a feature differs somewhat across RNA types, but significance alone is inflated by large sample size. We therefore report the rank-based effect size

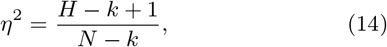

for which larger values indicate stronger between-type rank separation.

Applied directly to the sequence profiles, this effect size is confounded by sequence length. The pooling in Equation (12) is an average over tokens, so a feature that activates on a bounded number of positions has its profile value reduced roughly in proportion to sequence length, and RNA types in the holdout differ systematically in length (Supplementary Figure S1). A feature that responds to nothing but sequence length would therefore score a large *η*^2^, simply because the shortest RNA types are distinct types; *η*^2^ computed on the profiles cannot separate that case from genuine RNA-type selectivity. We therefore remove length before testing. Since the statistic is rank-based, we do so by rank-residualisation: for feature *i* we regress the ranks of its sequence profile on the ranks of sequence length,

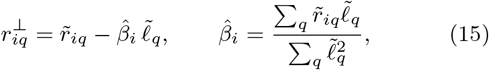

where 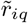 and 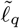 are the mean-centred ranks of the sequence profile *v*_*iq*_ and of the length of sequence *q*, and apply Equations (13) and (14) to the residuals 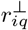. Every *η*^2^ and every RNA-type ranking reported below is computed this way, so it reflects the between-type separation that a monotone dependence on length cannot account for. The control is deliberately conservative: type-specific signal that happens to covary with length is removed along with the confound. Supplementary Figure S9 reports what the control removes. RNA-type association significance is corrected across tested SAE features using Bonferroni adjustment at *α* = 0.01, and is reported separately from *η*^2^ because very small *p*-values are expected at this holdout size.

For descriptive feature rankings, the preferred RNA type is the type with the largest mean activation. Fold enrichment is the preferred-type mean activation divided by the global mean activation, and selectivity is the preferred-type mean divided by the sum of the 16 unweighted type means. These descriptive quantities are computed on the raw activations, since the length residuals are not interpretable as activation magnitudes.

To test whether the sparse profiles are useful as representations, not merely statistically associated with labels, we run a simple *k*-nearest-neighbour probe with *k* = 7, cosine distance, and five-fold stratified cross-validation. The dense BiRNA-BERT baseline uses the analogous mean-pooled hidden states from the same layer and the same train/test splits. The SAE probe retains dimensions whose sequence-mean activation exceeds 0.05 in at least one holdout sequence. Because RNA types in this holdout differ systematically in length, we also run the identical probe on two sequence-only representations that use no model at all: the nucleotide length of each sequence, and its four mononucleotide frequencies. These establish how much of the RNA-type neighbourhood structure is recoverable from superficial sequence statistics, and therefore how much of the learned representations’ performance is attributable to the model. The primary score is balanced accuracy,

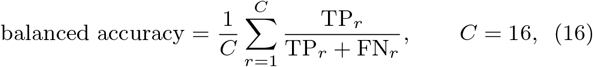

which is macro recall: every RNA type contributes equally, so the dominant class cannot hide poor performance on rare RNA types. For PCA visualisation, we sample at most 375 sequences per RNA type, which yields 4,417 sequences: eight types reach that cap and the remaining eight contribute every sequence they have. Dense profiles use all 768 mean-pooled hidden-state dimensions; SAE profiles retain active dimensions and, for the projection, the 256 dimensions with the largest variance. We report the variance explained by the first two components, silhouette score under the RNA-type labels, and adjusted Rand index (ARI) for a 16-cluster *k*-means solution in the two-dimensional projection.

## 3. Results and Discussion

### 3.1. Sparse Reconstruction Is Faithful Across Layers

Before assigning biological meaning to sparse features we first ask whether the SAEs faithfully preserve the original BiRNA-BERT state representations. Across each analysed depth the reconstruction fidelity is extremely high (Table 1). Explained variance remains above 0.99997 at every analysed depth, cosine similarity is effectively one, and sequence recovery stays near 99.7%. It also preserves the output distribution of the language model, with very small KL divergence and near zero cross entropy changes. The sparsity profile is also visible across layers: layer 0 uses 15.9% active dictionary features per token on average, while layers 5 and 11 are slightly sparser at 12.6% and 12.9%, respectively, while preserving reconstruction. These results mean the biological analyses below compare sparse representations that are similarly faithful across depth, rather than differences caused by distorted reconstruction.

### 3.2. Sparse Features Align with Secondary Structure

The structure-level analysis tests whether sparse features align with nucleotide-level bpRNA secondary-structure labels after BPE offset propagation.

The bpRNA-90 background is itself highly imbalanced (Figure 1(a)): Stem and Hairpin-loop labels are the two largest classes (46.4% and 19.6%, respectively). Pseudoknots are absent from this retained subset. This class imbalance makes raw selectivity scores difficult to interpret in isolation, as common structures are more likely to be the preferred label even when feature activations are randomly distributed. A sparse feature whose activations prefer Stem with selectivity around 0.46 therefore provides no evidence of Stem specificity from selectivity alone. High selectivity or high enrichment relative to the empirical background provides stronger evidence that the preferred label represents a specialised structural context.

**Figure 1.**
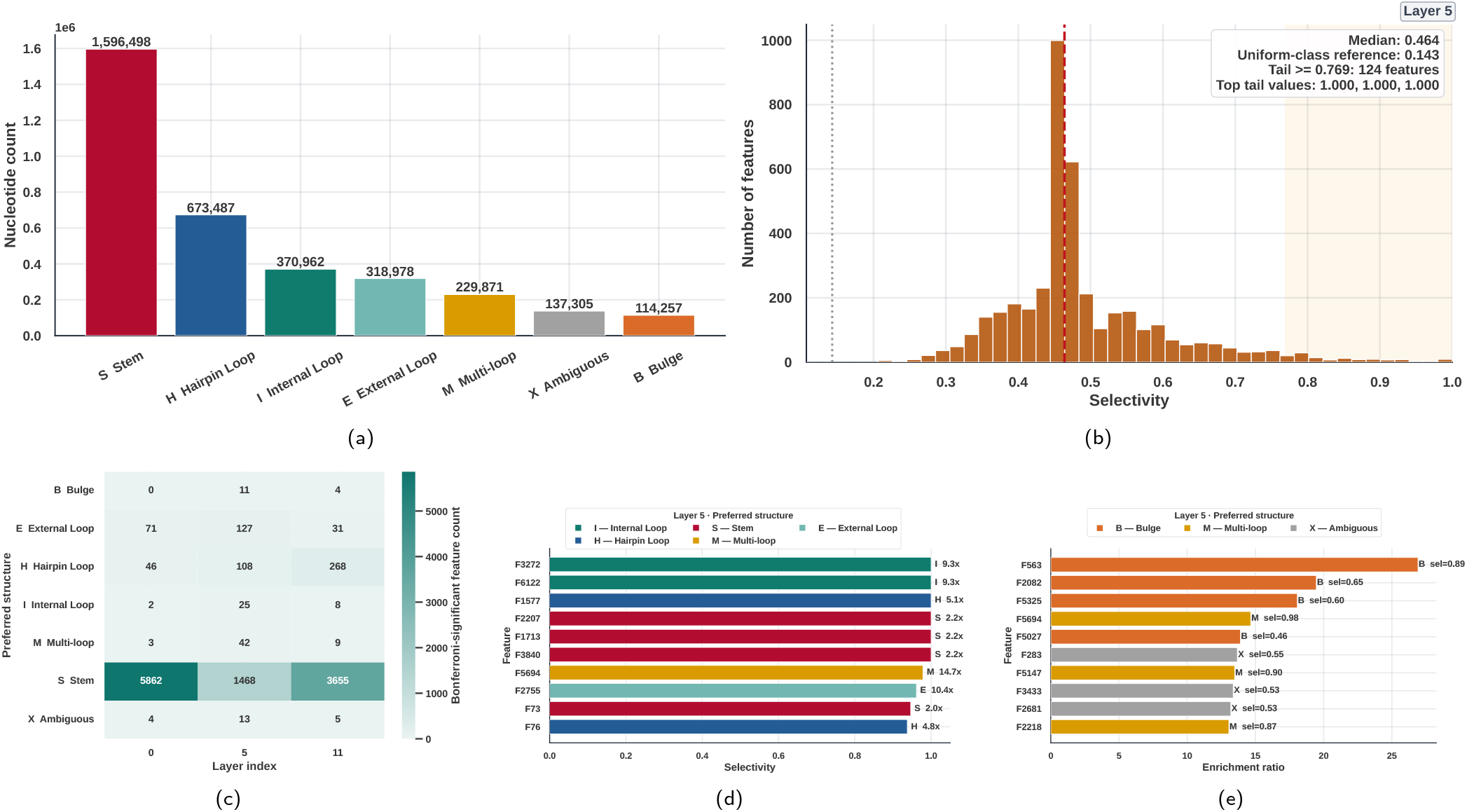
Secondary-structure alignment of sparse autoencoder (SAE) features with bpRNA-90 nucleotide secondary-structure labels. Figure (a) shows the background frequency of nucleotide secondary-structure classes in the evaluated bpRNA-90 subset. Figure (b) shows the distribution of layer-5 structural selectivity scores across evaluated SAE features; the dotted line marks the uniform-class reference 1*/*7. Figure (c) summarizes the number of statistically significant features assigned to each preferred structural label across layers. Figure (d) shows the most structure-selective layer-5 features, where bar lengths denote selectivity and annotations report the preferred structure class and enrichment ratio. Figure (e) shows the most structure-enriched layer-5 features, where bar lengths denote enrichment and annotations report the preferred structure class and selectivity score. Figures (d) and (e) are both restricted to Bonferroni-significant features.

Against that background, the layer-5 selectivity distribution (Figure 1(b)) has a median of 0.464, nearly identical to the medians at layers 0 and 11 (Supplementary Figure S2). Several layer-5 features reach the maximum selectivity value of 1.00, meaning 100% of their observed activations fall within one annotated structure type. Such perfectly selective features are missing from layer-0 representations. The distribution also shows a broader high-selectivity shoulder than layers 0 and 11, suggesting that middle-layer representations have shifted from broad sensitivity to more specific specialisation.

The cross-layer pattern in Table 2 reveals a trade-off between the number and strength of structure-associated features. Layer-0 achieves the highest absolute number of Bonferroni-significant features (5,988 of 6,139; 97.5%), but the mean enrichment is only 1.04*×*. Almost every layer-0 feature is statistically associated with some structure class, yet the associations are weak: activating a layer-0 feature barely shifts the probability distribution over structure labels beyond what the background already predicts. Layer 5 shows the opposite pattern: only 1,794 of 4,053 tested features (44.3%) pass Bonferroni correction, while mean enrichment across all tested features rises to 1.61*×*, indicating stronger concentration on preferred structural contexts overall. Layer 11 occupies an intermediate position (72.6% significant, mean enrichment 1.21*×*), suggesting that the final layer still carries structure-aligned features but with less enrichment than the layer-5 representations.

The class-count heatmap in Figure 1(c) adds an important qualification. Stem-selective features dominate at every layer. The dominance is most extreme at layer 0, with 5,862 of the 5,988 significant features assigned to Stem, while several non-Stem classes are represented by only a few features and no feature prefers Bulge. Despite the substantial drop in the number of significant features, layer 5 shows a less concentrated distribution on Stem. This shift is especially pronounced for Internal Loop and Multi-loop, whose absolute feature counts increase from 2 to 25 and from 3 to 42, respectively, together with 11 Bulge-preferring features. Stem dominance returns at layer 11, accompanied by an increase to 268 Hairpin-Loop-preferring features.

The strongest individual layer-5 features are shown from two angles in Figures 1(d) and 1(e). Both rankings are restricted to Bonferroni-significant features. Ranked by selectivity, the ten leading features span five structural classes (four prefer Stem, two Internal Loop, two Hairpin Loop, one Multi-loop, and one External Loop), and six of them are perfectly selective, firing only inside a single annotated structural context. Ranking by enrichment instead surfaces a largely disjoint set of features that prefer the rarer Bulge, Multi-loop, and Ambiguous labels, led by the Bulge-associated F563 at 26.9*×* enrichment and selectivity 0.894. At the boundary layers, peak enrichment is lower (19.5*×* and 20.0*×*) and no layer-0 feature prefers Bulge at all, so the rarest structural class is reached only at layers 5 and 11 (Supplementary Figure S4). Structure composition profiles showing the full multi-class activation breakdown for the top features, including the absence of perfect selectivity at Layer 0 and the regression to Stem dominance at Layer 11, are detailed in Supplementary Figure S13.

#### 3.3. Family-Level Features Capture Broader RNA-Type Signals

The RNA-type analysis asks a different question from the structure analysis. A secondary-structure feature is local and nucleotide-level; a sequence-level feature may reflect composition, length, motif usage, or structural tendency across an entire sequence. We mean-pool token activations into sequence profiles and analyse them on the holdout set. The holdout is heavily imbalanced (Figure 2(a)): lncRNA (14,602 sequences; 58.4%), rRNA (2,535; 10.1%), misc RNA (2,245; 9.0%), and tRNA (1,439; 5.8%) together account for 83.3% of the data, while the smallest retained type, ribozyme, contributes 91 sequences (0.36%). This makes balanced metrics and effect sizes essential.

**Figure 2.**
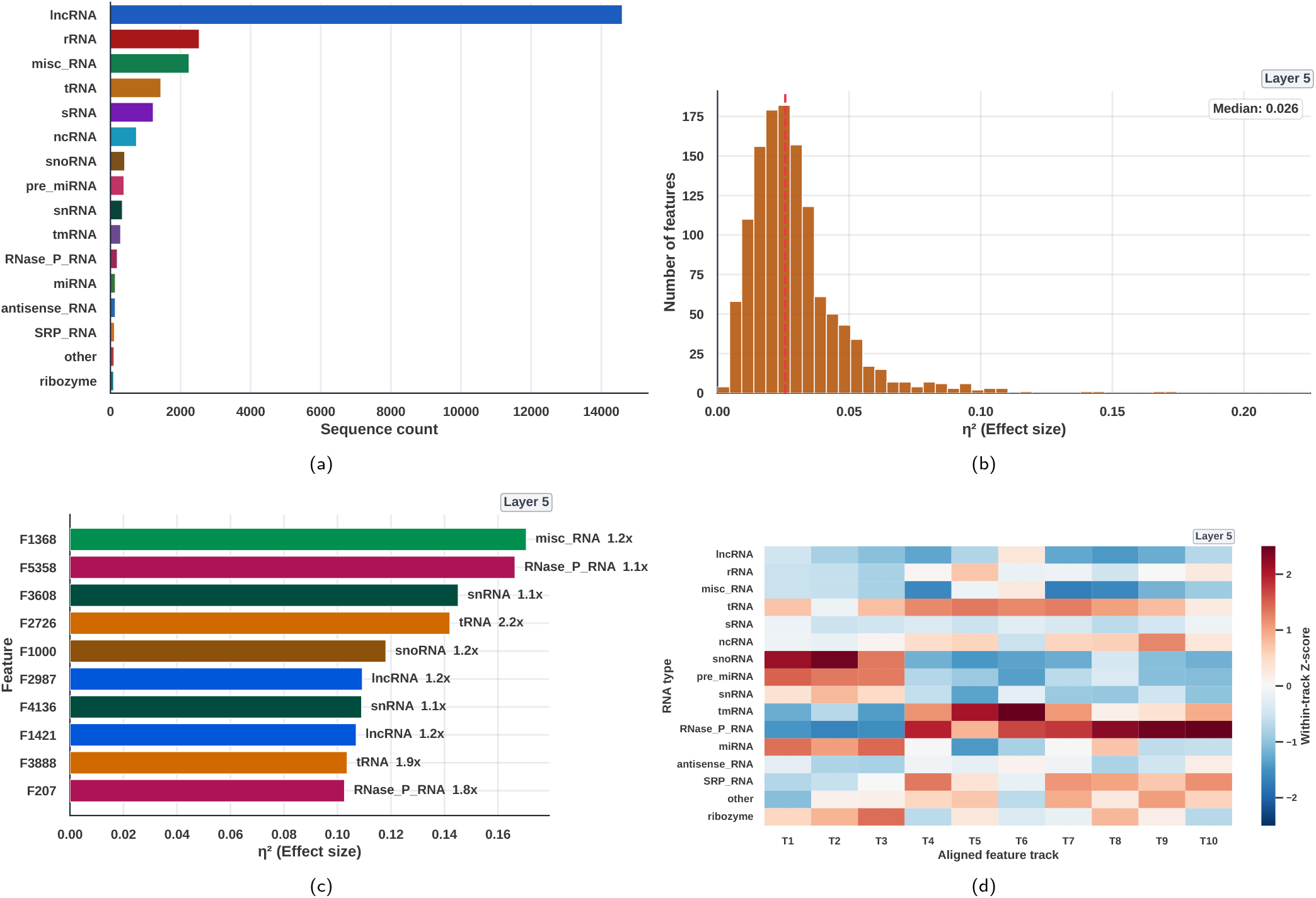
RNA-type alignment of layer-5 sparse autoencoder (SAE) sequence profiles on the similarity-filtered RNAcentral holdout (25,000 sequences, 16 RNA types). All *η*^2^ values are length-controlled (Section 2.4). Figure (a) shows the strongly imbalanced type composition, dominated by lncRNA, rRNA, and misc RNA. Figure (b) shows the distribution of Kruskal–Wallis effect sizes across the 1,237 eligible SAE features; the layer-5 median is *η*^2^ = 0.026 (dashed line), and the maximum is 0.171. Figure (c) ranks the ten features with the largest *η*^2^; annotations give each feature’s preferred RNA type and the ratio of its preferred-type mean activation to its global mean activation. The ten features prefer six different RNA types. Figure (d) shows the layer-5 members of ten feature tracks anchored at layer 0 and matched across layers by RNA-type profile cosine similarity. Colours are within-track Z-scores across RNA types; tracks T1–T3 peak on snoRNA, pre miRNA and miRNA, whereas T4–T10 peak on RNase P RNA and tmRNA.

The rank-based effect-size estimate *η*^2^ summarises between-type separation in a feature’s sequence-mean activation distribution, after the dependence of that distribution on sequence length has been removed (Section 2.4). The resulting distribution across the 1,237 eligible layer-5 features is right-skewed (Figure 2(b)), with median *η*^2^ = 0.026 and maximum *η*^2^ = 0.171. Every eligible feature passes Bonferroni correction at *α* = 0.01, yet the typical effect remains small. Cross-layer comparisons appear in Supplementary Figures S5–S7.

The features with the largest length-controlled effects are spread across RNA types (Figure 2(c)). The ten leading layer-5 features prefer six different types: misc RNA (F1368, *η*^2^ = 0.171), RNase P RNA (F5358, 0.166; F207, 0.103), snRNA (F3608, 0.145; F4136, 0.109), tRNA (F2726, 0.142; F3888, 0.104), snoRNA (F1000, 0.118), and lncRNA (F2987, 0.109; F1421, 0.107). Their fold enrichments range from 1.1 to 2.2*×* over each feature’s global mean activation. The two types carrying the most stereotyped sequence-and-structure constraints are represented here. tRNAs are constrained enough that covariance models built from exactly those constraints identify tRNA genes at near-perfect sensitivity and specificity (Chan et al., 2021; Phizicky and Hopper, 2010), and canonical snoRNAs are defined by class-specific C/D or H/ACA box motifs embedded in characteristic secondary structures (Jorjani et al., 2016). Neither type dominates the ranking, however.

Enrichment ranks features differently: the ten most enriched layer-5 features prefer only RNase P RNA and tRNA, yet reach *η*^2^ of just 0.02–0.10 and none appears among the ten above (Supplementary Figure S6). A feature can prefer its favoured type strongly relative to its own average activation while still separating the 16 types poorly, so we treat effect size as the primary ranking and enrichment as a descriptive companion.

The aligned heatmap (Figure 2(d)) places these features against all 16 RNA types. Columns T1–T10 are anchored by the ten highest-*η*^2^ layer-0 features and matched one-to-one at other layers by cosine similarity between their 16-type mean-activation profiles; cell colour is the within-track Z-score. The tracks separate into two groups that respond to different parts of the type vocabulary. T1–T3, anchored by miRNA-preferring layer-0 features, are strongly positive for snoRNA, pre miRNA, and miRNA and negative for RNase P RNA and tmRNA; T4–T10, anchored by tRNA-, tmRNA-, and RNase P RNA-preferring features, invert that pattern, peaking on RNase P RNA and tmRNA. Individual tracks are therefore selective for particular groups of types, and the alignment is preserved across depth (Supplementary Figure S8).

### 3.4. Sparse Profiles Retain and Improve Useful RNA-Type Neighbourhoods

We next asked whether sparse profiles are useful as representations, not merely statistically associated with labels. A deliberately simple *k*NN probe (*k* = 7, cosine distance, five-fold stratified cross-validation) tests whether sequences from the same RNA type become neighbours. Layer 5 is the best setting overall, and there the SAE raises balanced accuracy from 0.328 for the dense BiRNA-BERT embedding to 0.359 (Figure 3(a), Table 3). Layer 11 also improves slightly (0.284 vs. 0.275) whereas layer 0 declines (0.250 vs. 0.258), so sparse profiles improve RNA-type neighbourhoods at the middle and final layers but not uniformly across depth. Balanced accuracy is macro recall, which is the informative metric here because every type contributes equally despite the holdout imbalance.

**Table 3.** *k*-nearest-neighbour (*k*NN) RNA-type classification on mean-pooled sequence representations from the 16-type RNAcentral holdout (*k* = 7, cosine distance, five-fold stratified cross-validation). Balanced accuracy is macro recall, and macro precision is the unweighted mean of per-class precision values. The sequence-only baselines use the same protocol and are layer-independent: length is the nucleotide length after preprocessing, composition is the four mononucleotide frequencies. A one-dimensional profile is degenerate under cosine distance, since all such vectors are collinear; evaluated with Euclidean distance the length-only baseline reaches 0.171 balanced accuracy.

| Layer | Representation | Macro precision | Balanced acc. |
| --- | --- | --- | --- |
| – | Length | 0.037 | 0.062 |
|  | Composition | 0.148 | 0.102 |
|  | Length + comp. | 0.352 | 0.298 |
| 0 | BiRNA-BERT | 0.411 | **0.258** |
|  | SAE | **0.452** | 0.250 |
| 5 | BiRNA-BERT | 0.489 | 0.328 |
|  | SAE | **0.514** | **0.359** |
| 11 | BiRNA-BERT | 0.398 | 0.275 |
|  | SAE | **0.400** | **0.284** |

**Figure 3.**
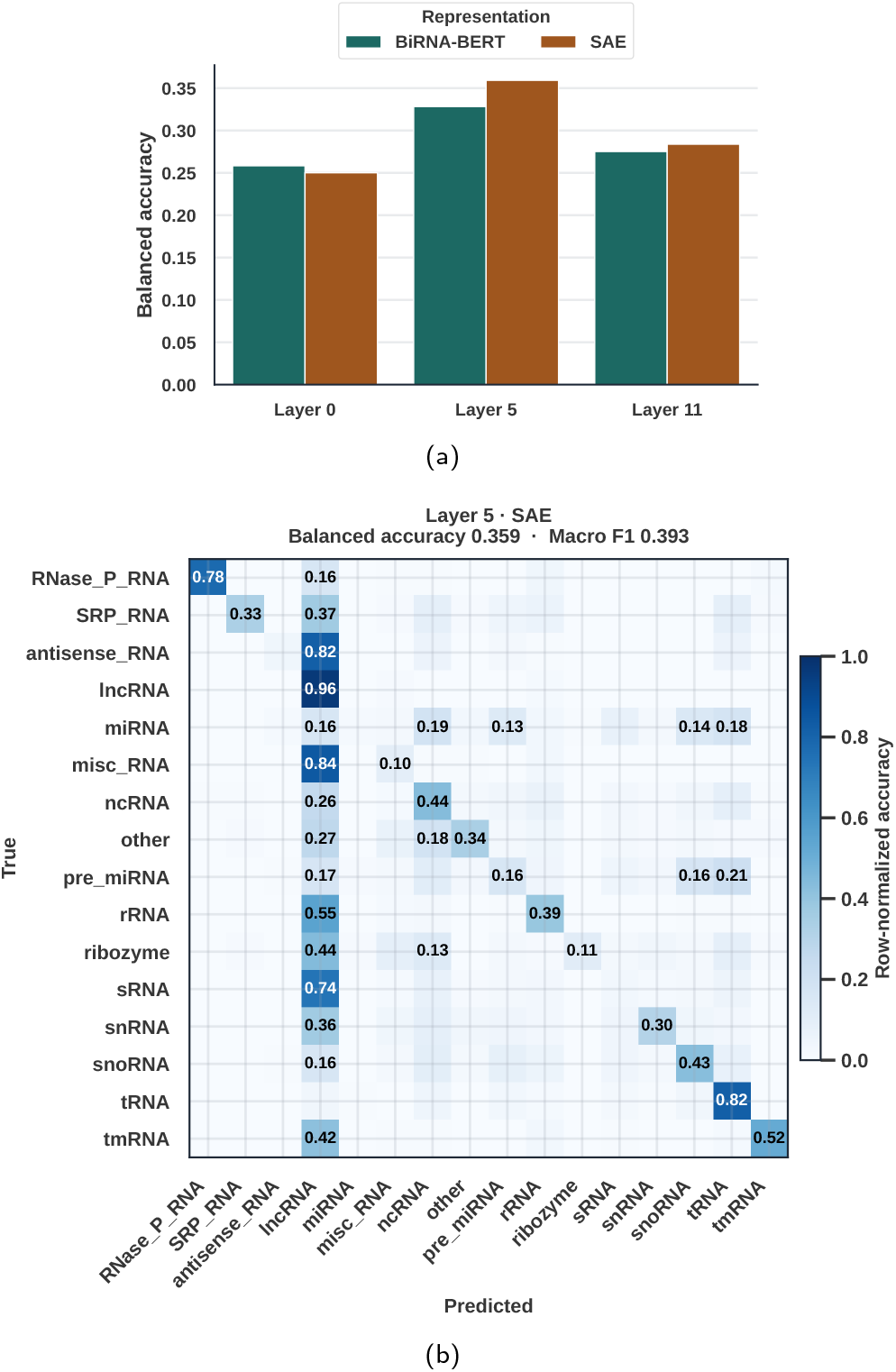
*k*-nearest-neighbour (*k*NN) RNA-type evaluation on mean-pooled sequence representations from the 16-type RNAcentral holdout (*k* = 7, cosine distance, five-fold stratified cross-validation). Figure (a) compares balanced accuracy for dense BiRNA-BERT embeddings and SAE profiles across layers, and Figure (b) shows the layer-5 SAE confusion matrix. Balanced accuracy is macro recall, which is the more informative metric for this strongly imbalanced holdout.

Because RNA types in this holdout separate strongly by length, a probe of this kind could in principle succeed on superficial sequence statistics alone. The effect-size analysis mentioned earlier guards against this at the level of individual features, but the *k*NN probe uses the whole profile and needs its own control. We therefore ran the identical protocol on two hand-built representations: sequence length, and mononucleotide composition (Table 3). Neither is competitive on its own, but together they reach 0.298 balanced accuracy, recovering a substantial share of what the learned representations achieve and more than either BiRNA-BERT or the SAE manages at layers 0 and 11. The layer-5 representations nonetheless stay clearly ahead (0.328 and 0.359), so the middle-layer neighbourhood structure is not merely a restatement of length and composition.

The class-level behaviour of the layer-5 SAE (Figure 3(b)) shows that the errors are structured rather than diffuse, and that their structure follows abundance within a length regime. Among the long RNA types, lncRNA acts as a sink: it recovers almost perfectly (0.96) while misc RNA (0.84), antisense RNA (0.82), sRNA (0.74), rRNA (0.55), and tmRNA (0.42) are predominantly assigned to it. Among the short types the same thing happens at a different address, with miRNA and pre miRNA drawn towards tRNA (0.18 and 0.21) and snoRNA (0.14 and 0.16). In both regimes the misassignments flow towards the most abundant type of comparable length, since lncRNA holds 58.4% of the holdout and tRNA is the most abundant short type. Only types with both distinctive folds and adequate support hold their own diagonal, namely tRNA (0.82), RNase P RNA (0.78), and tmRNA (0.52). We note that biological heterogeneity may compound the lncRNA sink, since lncRNA is an operationally broad category rather than a structurally coherent family (Cabili et al., 2011; Statello et al., 2021); but heterogeneity and the class prior are confounded in this holdout, and the present analysis cannot separate them.

The dense BiRNA-BERT matrix for the same layer (Supplementary Figure S11) locates the SAE’s advantage.

The lncRNA sink and the dominant diagonals are unchanged (lncRNA 0.96 under both, tRNA 0.83 and 0.82); the gain lies entirely in minority types such as snRNA (0.21 to 0.30), snoRNA (0.34 to 0.43), and SRP RNA (0.23 to 0.33), which is precisely what moves macro recall. Extended *k*NN results for all layers appear in Supplementary Figures S10 and S12.

### 3.5. Principal Component Analysis Provides Diagnostic Evidence

We perform principal component analysis (PCA) on the layer-5 representations to visualise the RNA-type separation of dense BiRNA-BERT embeddings and SAE profiles. In

Figure 4, the first two principal components of the layer 5 representations explain 48.0% of variance for dense BiRNA-BERT embeddings and 54.2% for the variance-selected SAE profiles. Both projections show broad RNA-type groupings, but many types remain overlapping. The SAE projection has a slightly higher ARI (0.095 vs. 0.085), while the silhouette scores are indistinguishable (−0.197 for both). The negative silhouette scores indicate that, on average, within-type cohesion is weaker than separation from at least one neighbouring RNA type.

**Figure 4.**
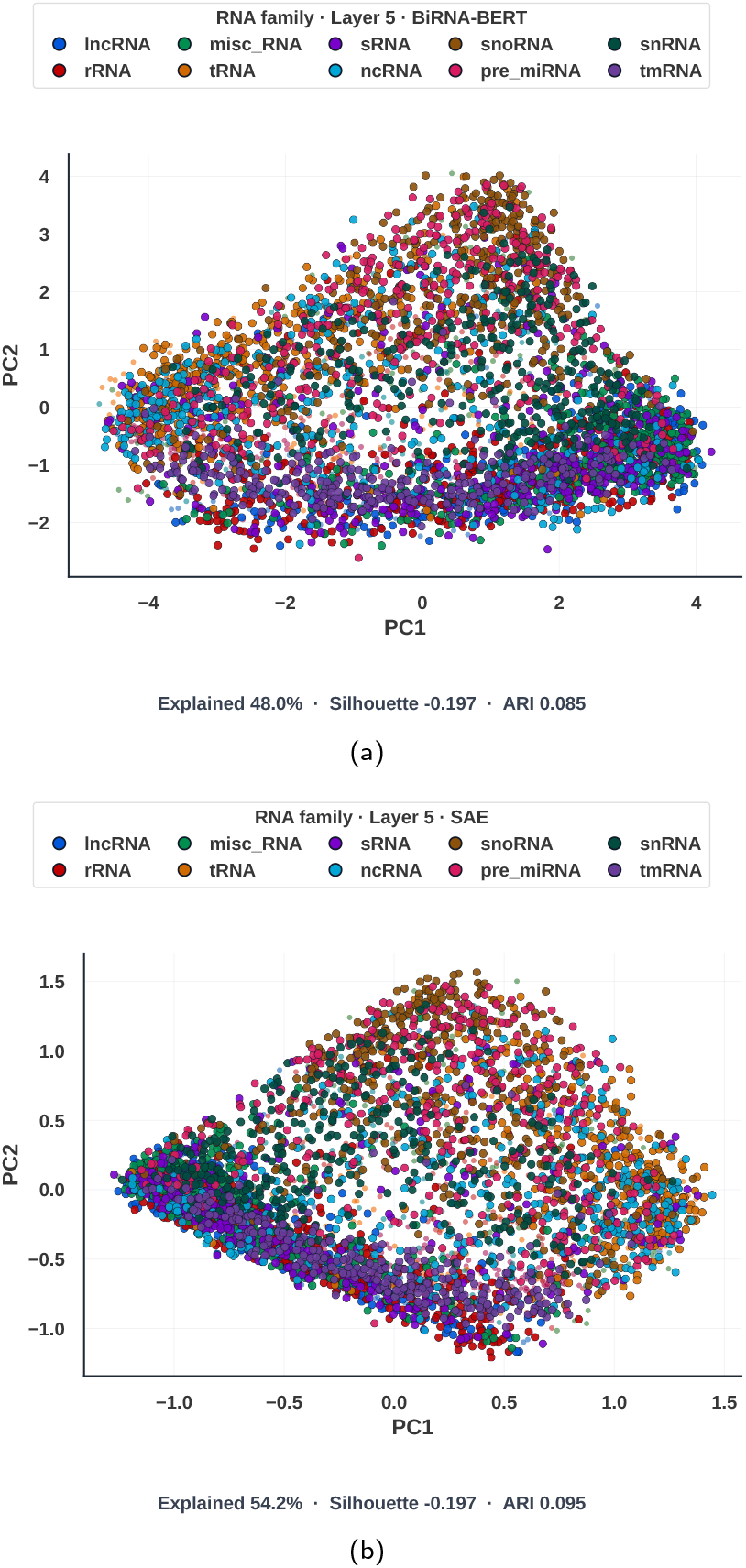
Two-dimensional principal component analysis (PCA) of layer-5 sequence representations coloured by RNA type. Each point is one of 4,417 sequences drawn from the 16-type RNAcentral holdout by sampling at most 375 per type (Section 2.4). The legend shows the 10 most abundant types. Figure (a) and Figure (b) compare dense BiRNA-BERT embeddings and SAE profiles; explained variance, Adjusted Rand Index (ARI), and silhouette score summarize how well the projection separates RNA types.

Interestingly, the length confound identified above also governs the leading components: in the layer-5 sample, sequence length correlates with the SAE projection at *r* = −0.65 on PC1 and −0.57 on PC2, and with the dense BiRNA-BERT projection at *r* = 0.42 and −0.74. Much of the visible organisation in both panels therefore orders sequences by length rather than by type.

## 4. Conclusion

SPIRAL shows that sparse autoencoders can turn dense BiRNA-BERT hidden states into a more inspectable feature space while preserving the behaviour of the underlying language model. Unlike most prior RNA representation studies, which primarily evaluate downstream predictive performance, our analysis asks what biological signals are exposed by the representation itself. We train layer-wise SAE dictionaries, verify that reconstruction remains faithful, and then connect individual sparse features to nucleotide-resolution secondary structure, RNA-type identity, and sequence-neighbourhood geometry. This makes SPIRAL a useful mechanistic analysis framework.

We show that SAE features can be evaluated as biological objects rather than only as reconstruction units. At the local scale, features include high-selectivity and high-enrichment associations with secondary-structure contexts, including rarer classes such as Bulge and Multi-loop. At the sequence scale, mean-pooled SAE profiles reveal RNA-type-associated features and modestly improve balanced *k*NN accuracy over dense BiRNA-BERT embeddings on the holdout, staying ahead of length- and composition-only baselines. The cross-layer comparison adds a further distinction from prior single-layer analyses: early and late layers contain many statistical structure associations, but the middle layer has the largest mean structural enrichment.

Measuring these features also requires care about what mean pooling encodes. RNA types separate strongly by sequence length, and a token-mean profile makes any feature with bounded activation length-sensitive by construction, so an uncontrolled effect size rewards features that do no more than detect short sequences. We control for length in feature-level association testing and explicitly quantify its contribution to the sequence-level kNN and PCA analyses. Doing so changes which features rank highest, replacing an apparent population of tRNA detectors with a smaller and more type-diverse set. Tokenizer-aware offset propagation is another practical component of this analysis, allowing token-level activations to be compared with nucleotide-level bpRNA labels without collapsing each token to a single position.

These results show that SAEs expose existing biological structure in BiRNA-BERT. Future work can build directly on this feature space by fine-tuning SAE embeddings for downstream RNA tasks, comparing dictionary size and sparsity settings across RNA language models, and testing whether validated sparse features can be used as control handles for steering or designing sequences with intended family, motif, or structural properties.

## Use of generative AI

Generative AI tools were used as writing and coding aids only: language editing (grammar, wording and clarity) of author-written text, and assistance with implementation and refactoring of the analysis code.

## Data availability

The analyses use public RNAcentral and bpRNA-90 data. The research code is freely available to non-commercial users under the MIT license at https://github.com/SadatHossain01/SPIRAL. The same code, together with the filtered RNAcentral holdout, the processed bpRNA evaluation subset, the trained sparse-autoencoder checkpoints, configurations, and reproduction instructions, is archived on Zenodo at https://doi.org/10.5281/zenodo.21891845 (Hossain et al., 2026).

## Author contributions

M.S.H. and M.R.S. designed and implemented the analyses, trained sparse autoencoders, generated figures and drafted the manuscript. M.T.T. proposed the initial SAE direction and provided methodological feedback. M.S.R. supervised the project. All authors reviewed and edited the manuscript.

1 https://huggingface.co/buetnlpbio/birna-bert

2 https://huggingface.co/datasets/multimolecule/rnacentral

3 https://huggingface.co/datasets/multimolecule/bprna-90

