## Supplementary File for "Sparse Autoencoders Reveal Structural and Family-level Features in BiRNA-BERT"

### Supplementary Material for “Sparse Autoencoders Reveal Structural and Family-level Features in BiRNA-BERT”

#### Abstract

**This supplementary material provides additional layer-wise structure and RNA-family analyses, holdout construction details, and extended kNN results supporting the SPIRAL manuscript.**

**Table S1.** Holdout construction statistics with `cd-hit-est-2d`.

| Step | Count |
|---|---|
| Training-reference sequences | 1,000,000 |
| Candidates scanned from RNAcentral stream | 152,000 |
| Unique sequences retained after similarity filtering | 25,424 |
| Eligible after minimum RNA-type count | 25,347 |
| Eligible RNA types | 16 |
| Retained in final holdout | **25,000** |

**Table S2.** RNA-type distribution in the RNAcentral holdout.

| RNA type | Count | % |
|---|---|---|
| lncRNA | 14,602 | 58.41 |
| rRNA | 2,535 | 10.14 |
| misc_RNA | 2,245 | 8.98 |
| tRNA | 1,439 | 5.76 |
| sRNA | 1,222 | 4.89 |
| ncRNA | 744 | 2.98 |
| snoRNA | 408 | 1.63 |
| pre_miRNA | 388 | 1.55 |
| snRNA | 341 | 1.36 |
| tmRNA | 292 | 1.17 |
| RNase_P_RNA | 196 | 0.78 |
| miRNA | 140 | 0.56 |
| antisense_RNA | 139 | 0.56 |
| SRP_RNA | 115 | 0.46 |
| other | 103 | 0.41 |
| ribozyme | 91 | 0.36 |
| **Total** | **25,000** | **100.00** |

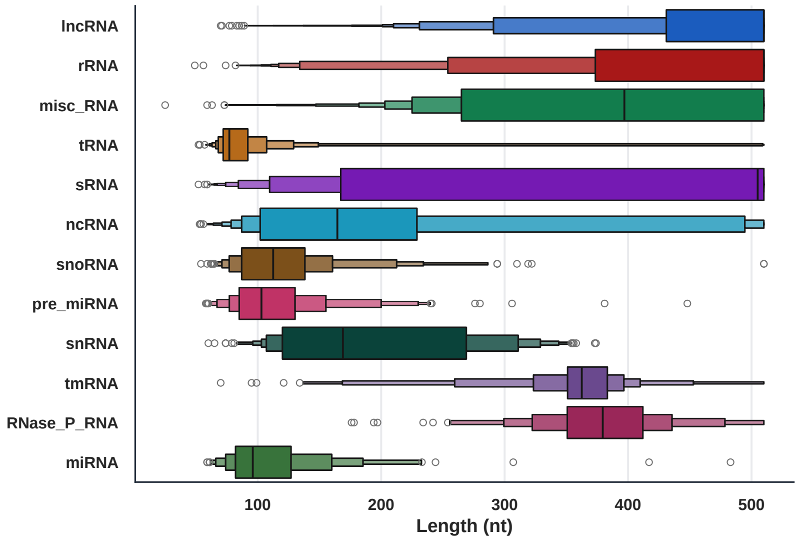


**Figure S1** Sequence-length distribution by RNA type in the 25,000-sequence RNAcentral holdout, ordered by frequency. Sequences were truncated at 510 nt during preprocessing, so the longest types accumulate at that bound. Types separate strongly by length: tRNA, miRNA, pre_miRNA, snoRNA, and ribozyme have median lengths of 77–125 nt, whereas tmRNA, RNase_P_RNA, misc_RNA, sRNA, lncRNA, antisense_RNA, and rRNA have medians of 362 nt or more.

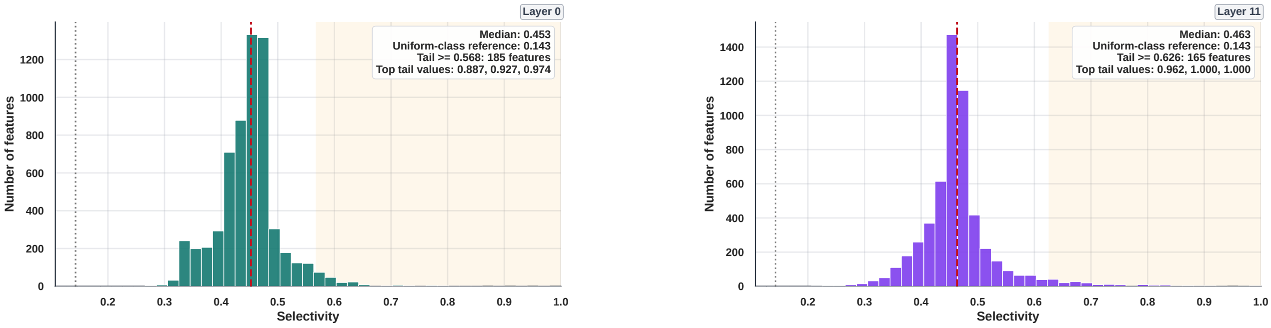


(a) (b)

**Figure S2** Structure selectivity distributions for layers 0 and 11. Dashed red lines mark the medians and dotted black lines mark the uniform-class reference ($1/7 \approx 0.143$). Both layers concentrate near selectivity $0.45$–$0.46$, with sparse high-selectivity tails.

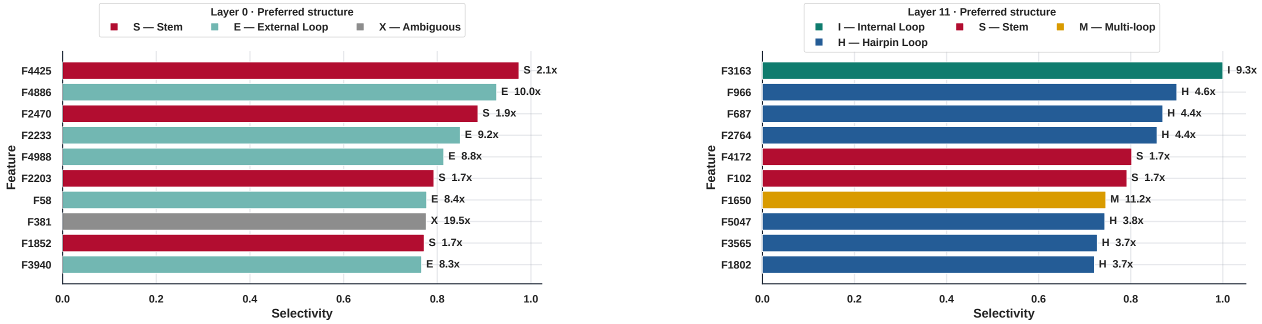


(a) (b)

**Figure S3** Top 10 structure-selective features at layers 0 and 11, restricted to Bonferroni-significant features. Bar length encodes selectivity, labels report enrichment, and colours indicate the preferred structure class. Layer 0 is split between External Loop (five features) and Stem (four), plus one Ambiguous feature, and reaches a maximum selectivity of 0.974. Layer 11 is instead dominated by Hairpin Loop (six features), with two Stem features and one each for Internal Loop and Multi-loop. Neither boundary layer reproduces the perfectly selective Internal Loop and Stem features seen at layer 5, apart from a single Internal Loop feature at layer 11.

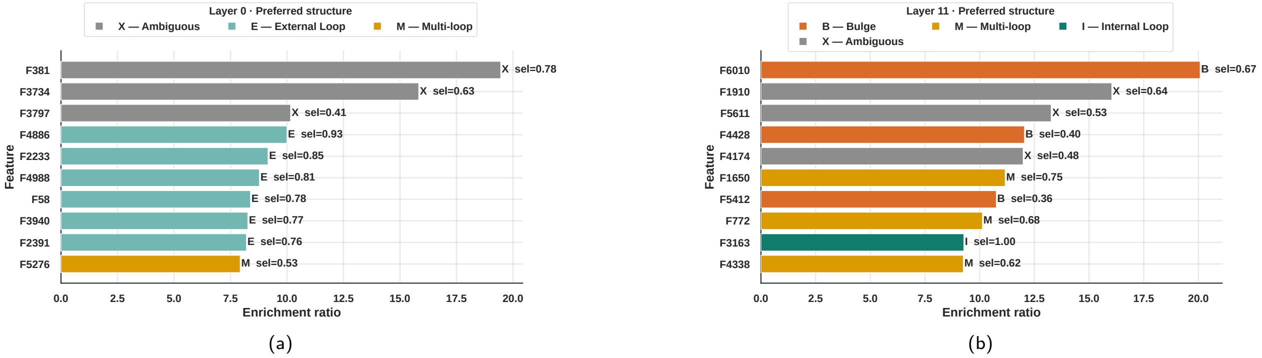


**Figure S4** Top 10 structure-enriched features at layers 0 and 11, restricted to Bonferroni-significant features, complementing the layer-5 ranking in the main text. Bar length encodes enrichment over the bpRNA background, labels report selectivity, and colours indicate the preferred structure class. Both boundary layers reach lower peak enrichment than layer 5 (19.5× at layer 0 and 20.0× at layer 11, against 26.9× at layer 5). The two layers also differ in which rare classes they reach: layer 0 contains no Bulge-preferring feature at all and its ranking is filled by Ambiguous and External Loop features, whereas layer 11, like layer 5, does contain Bulge-preferring features.

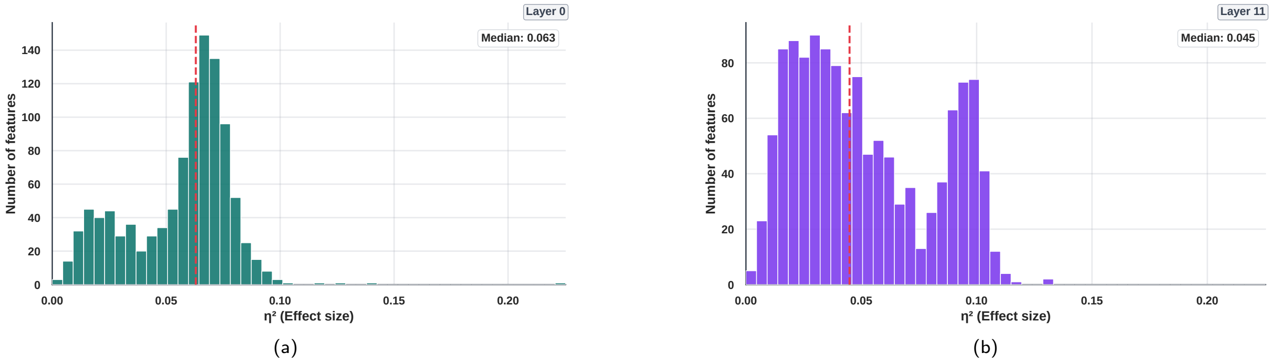


**Figure S5** Length-controlled Kruskal-Wallis $\eta^2$ distributions for layers 0 and 11, complementing the layer 5 distribution in the main text. Layer 0 has a dominant mode just above its median of $0.063$, whereas layer 11 remains visibly bimodal after the length control, with a second population of features near $\eta^2 \approx 0.095$ separated from the bulk; its median is $0.045$.

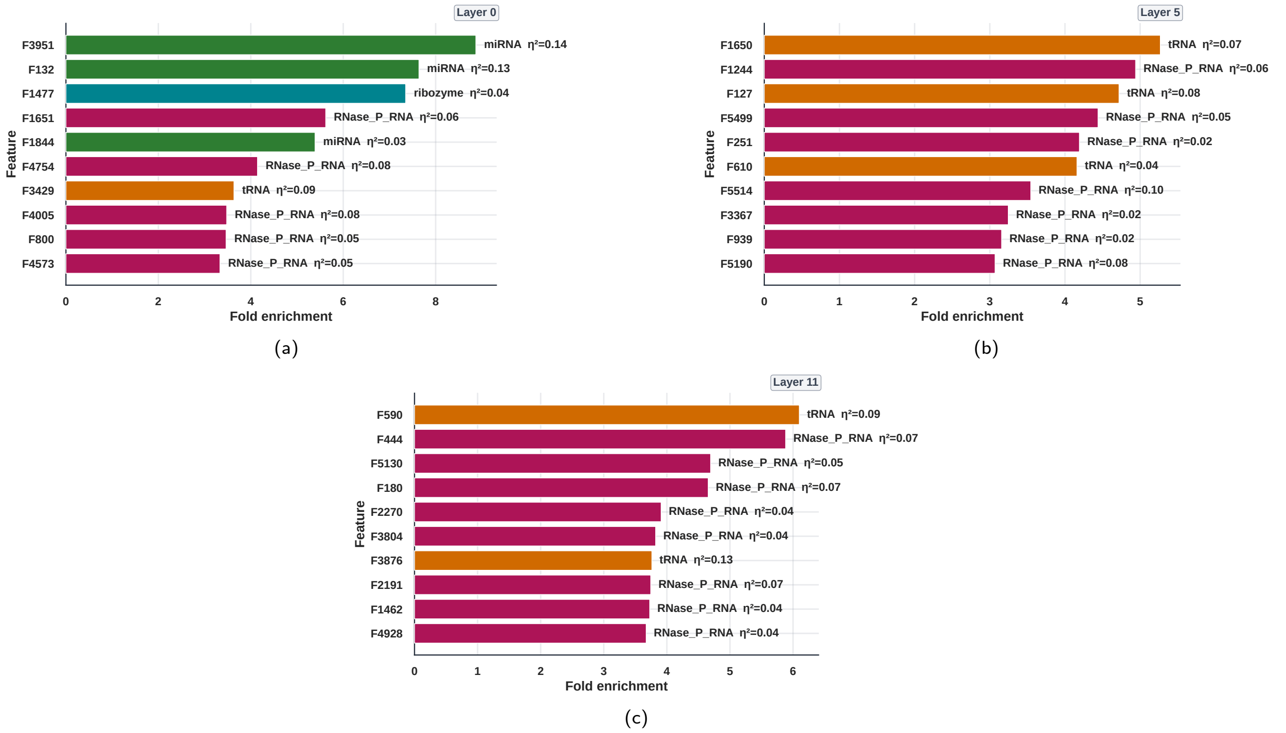


**Figure S6** Top 10 RNA-type-enriched features at all three layers. Bar length encodes preferred-type mean activation divided by global mean activation, colours indicate the preferred RNA type, and labels report each feature's length-controlled $\eta^2$. Layer 0 contains miRNA, ribozyme, RNase_P_RNA, and tRNA preferences, whereas layers 5 and 11 are dominated by RNase_P_RNA and tRNA. Enrichment and effect size rank features differently: the most enriched layer-5 features reach 3.1–5.3× but carry length-controlled effect sizes of only $\eta^2 = 0.02–0.10$, none of which places them among the ten features with the largest effect size. Enrichment measures how much a feature prefers its favoured type relative to its own average activation, which a feature can achieve while still separating the types poorly overall.

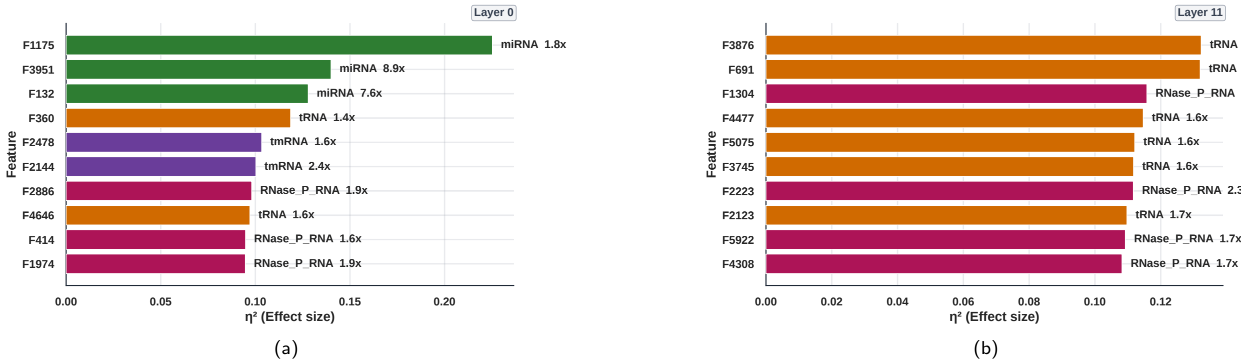


**Figure S7** Top 10 RNA-type-selective features ranked by length-controlled effect size at layers 0 and 11. Colours indicate preferred RNA type. Layer 0 spans four types (three miRNA, three RNase_P_RNA, two tRNA and two tmRNA features), whereas layer 11 concentrates on two types, with six tRNA-preferring and four RNase_P_RNA-preferring features.

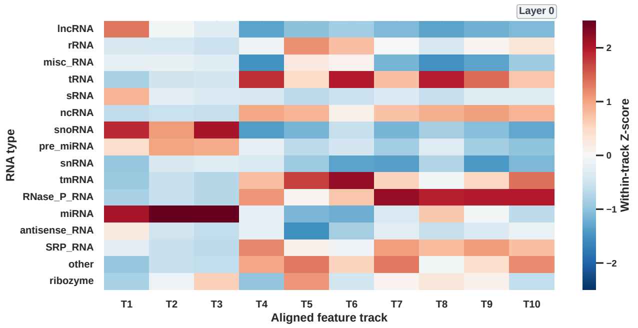


(a)

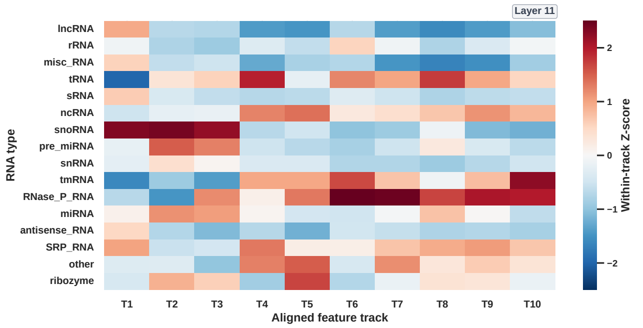


(b)

**Figure S8** Aligned mean-activation heatmaps for layers 0 and 11. Rows are the 16 RNA types ordered by holdout frequency. Columns are ten feature tracks anchored by the highest-$\eta^2$ layer-0 features and matched one-to-one at other layers by cosine similarity of their RNA-type mean activation profiles. Colours are within-track Z-scores across RNA types, so they show relative rather than absolute activation. Anchors are selected by length-controlled $\eta^2$: T1–T3 are miRNA-preferring, T4 and T8 are tRNA-preferring, T5 and T6 are tmRNA-preferring, and T7, T9 and T10 are RNase_P_RNA-preferring. The two-group organization described for layer 5 in the main text is preserved at both boundary layers.

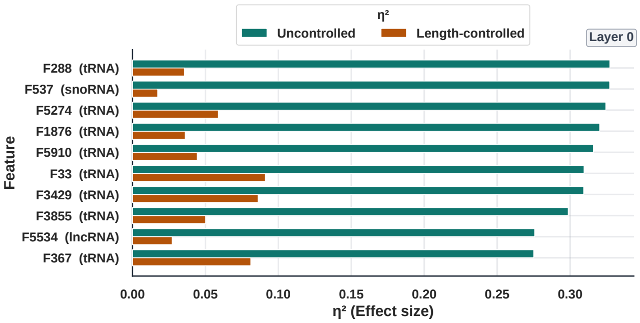


(a)

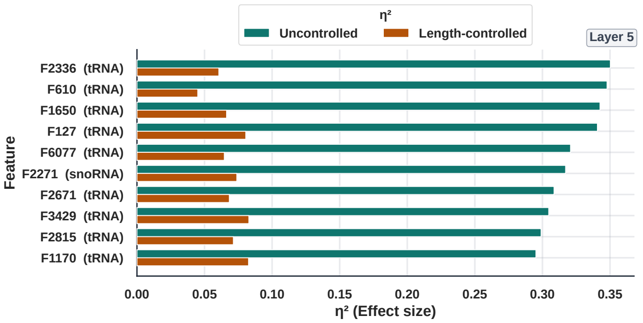


(b)

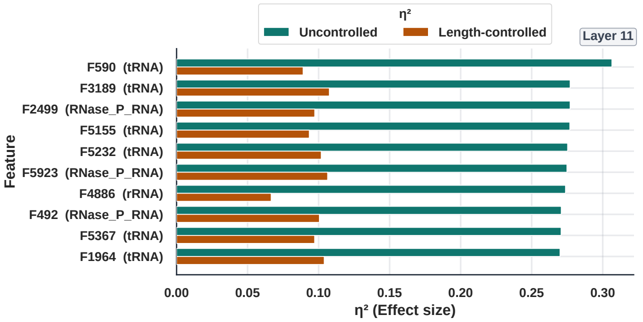


(c)

**Figure S9** The length control removal effect. Length-controlled RNA-type effect sizes have been reported throughout; this figure compares them against the uncontrolled values. Bars show, for each layer, the ten features with the largest *uncontrolled* $\eta^2$, together with the length-controlled $\eta^2$ of the same features. These features are strongly length-dependent and lose most of their effect once length is removed, which is why the uncontrolled ranking can be misleading: at layer 5 it is headed almost entirely by tRNA-preferring features, tRNA being the shortest type in the holdout. The confound affects all three layers and is strongest at layer 11; across all eligible features, the median $\eta^2$ retained after control is 75% at layer 0, 62% at layer 5, and 43% at layer 11.

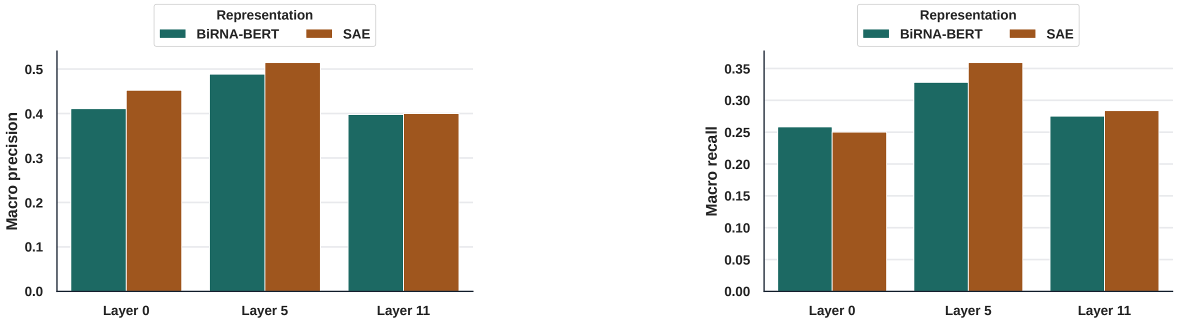


(a) (b)

**Figure S10** kNN macro precision and macro recall for dense BiRNA-BERT and SAE representations at layers 0, 5, and 11 ($k = 7$, cosine distance, 5-fold cross-validation). SAE representations improve macro precision at all three layers and macro recall at layers 5 and 11, but slightly reduce macro recall at layer 0.

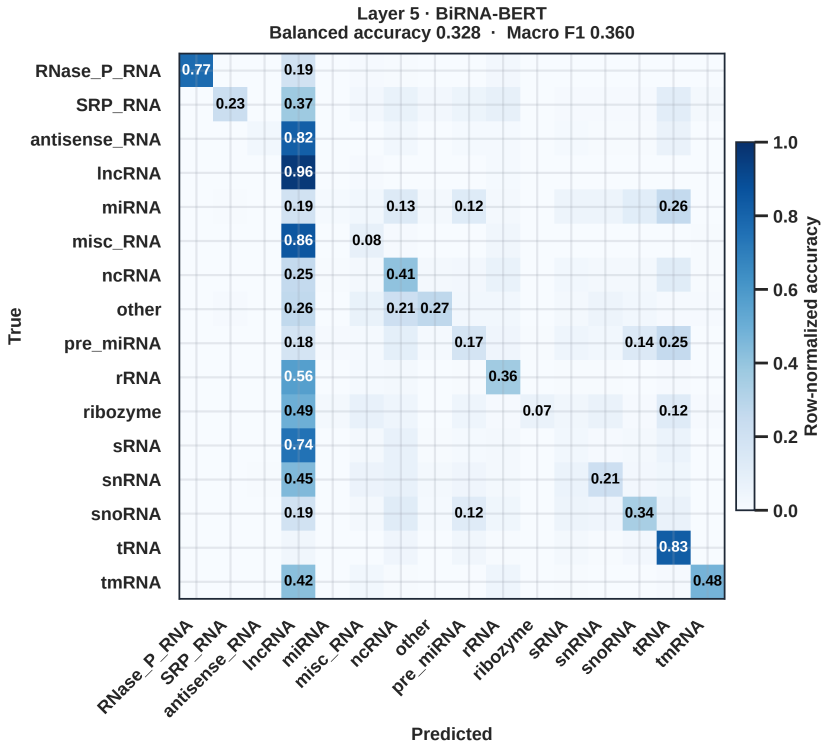


**Figure S11** kNN confusion matrix for the dense BiRNA-BERT representation at layer 5, the direct counterpart of the layer-5 SAE matrix in the main text. Comparing the two shows where the SAE's higher balanced accuracy (0.359 against 0.328) comes from. The two dominant diagonals are unchanged, with lncRNA at 0.96 under both representations and tRNA at 0.83 and 0.82; the gain is concentrated in the minority types, where the SAE raises recall for snRNA (0.21 to 0.30), snoRNA (0.34 to 0.43), SRP_RNA (0.23 to 0.33), other (0.27 to 0.34), tmRNA (0.48 to 0.52), ncRNA (0.41 to 0.44), and rRNA (0.36 to 0.39). Because balanced accuracy is macro recall, improvements confined to rare types move it even though the lncRNA sink persists in both representations.

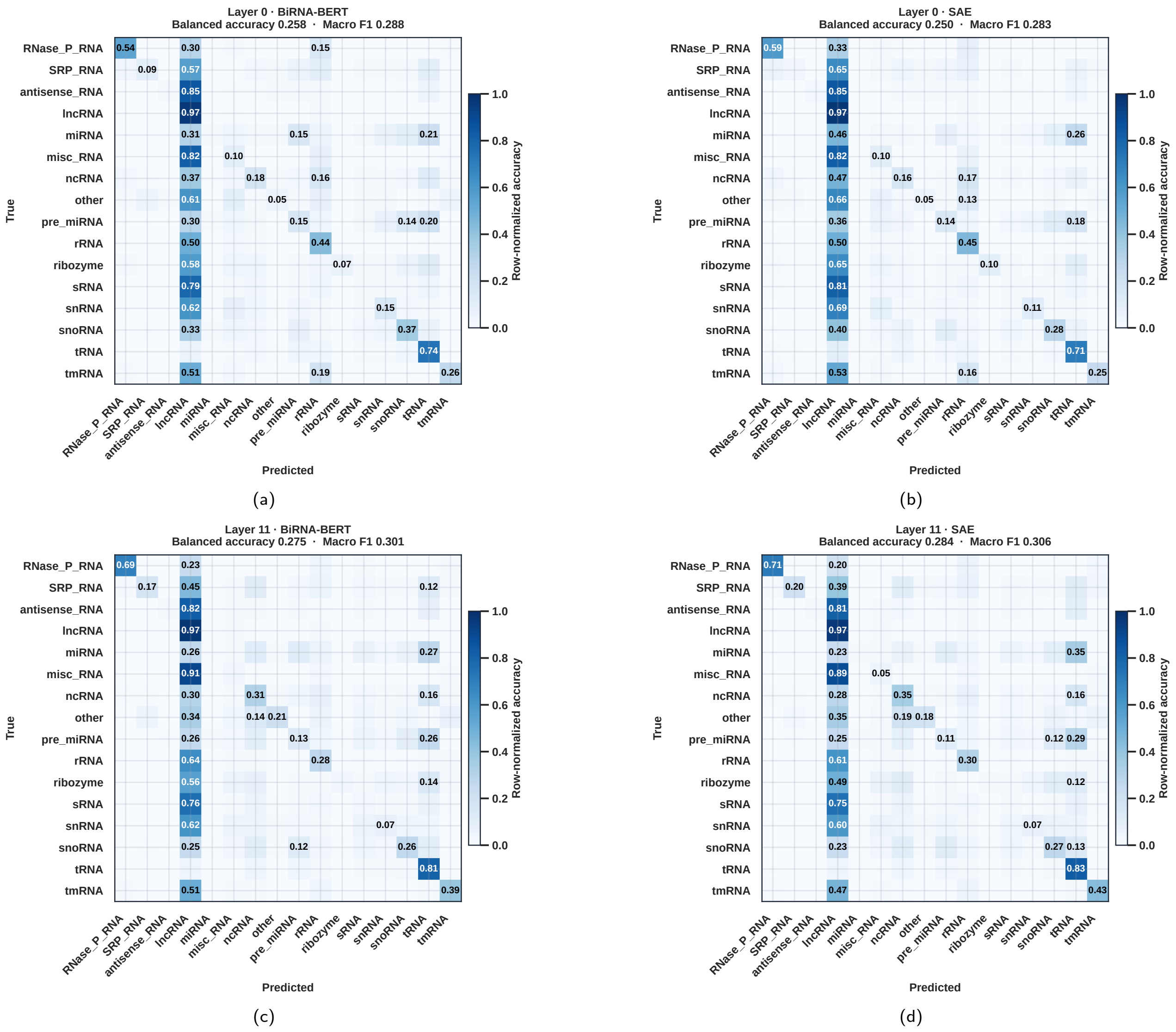


**Figure S12** kNN confusion matrices for boundary-layer representations. Figure (a) and Figure (b) show layer 0 (balanced accuracy: BiRNA-BERT $0.258$, SAE $0.250$), and Figure (c) and Figure (d) show layer 11 (BiRNA-BERT $0.275$, SAE $0.284$). lncRNA, miscRNA and tRNA have the strongest diagonal recall values, while many less frequent RNA types are predominantly predicted as lncRNA.

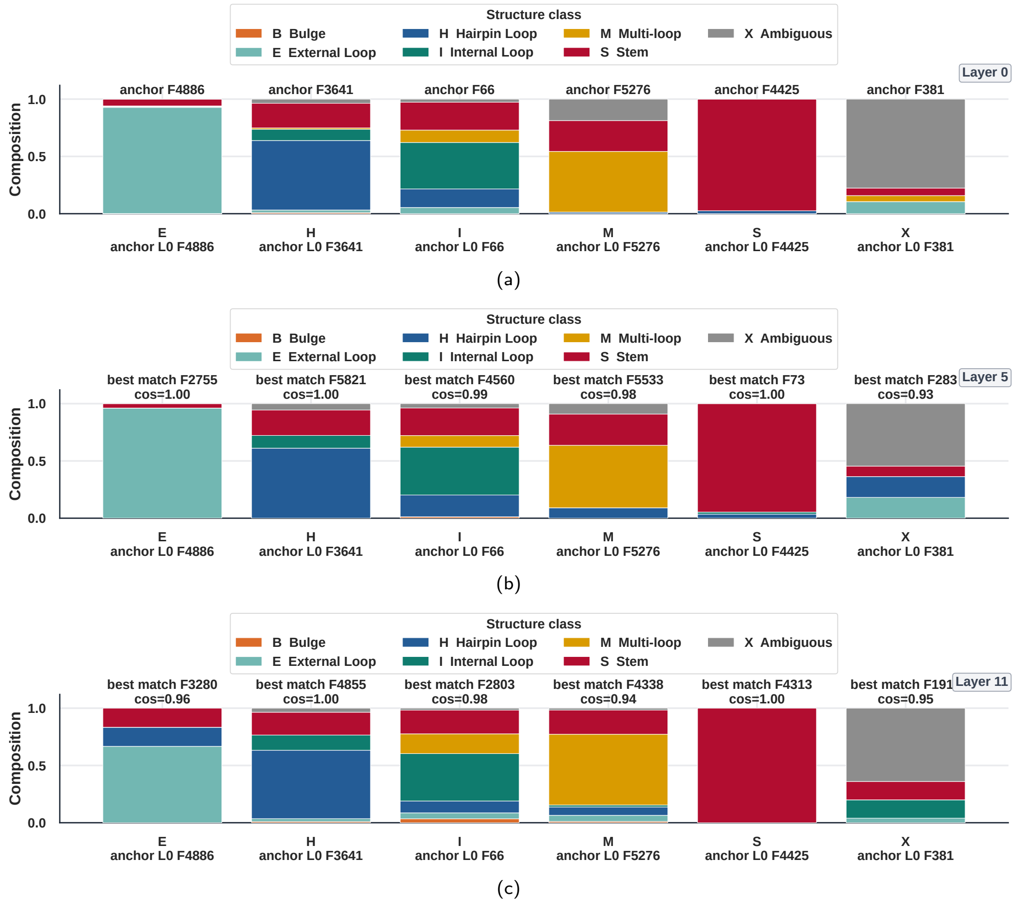


**Figure S13** Structure composition profiles for matched structure-selective features. Layer 0 anchor columns correspond to External Loop, Hairpin Loop, Internal Loop, Multi-loop, Stem, and Ambiguous preferences; layers 5 and 11 show the closest cosine-matched profile for each anchor. Stacked colours give the fraction of activations assigned to each structure class.
